# Long-read Sequencing Uncovers a Complex Transcriptome Topology in Varicella Zoster Virus

**DOI:** 10.1101/399048

**Authors:** István Prazsák, Norbert Moldován, Dóra Tombácz, Klára Megyeri, Attila Szűcs, Zsolt Csabai, Zsolt Boldogkői

## Abstract

**Background:** Varicella zoster virus (VZV) is a human pathogenic alphaherpesvirus harboring a relatively large DNA molecule. The VZV transcriptome has already been analyzed by microarray and short-read sequencing analyses. However, both approaches have substantial limitations when used for structural characterization of transcript isoforms, even if supplemented with primer extension or other techniques. Among others, they are inefficient in distinguishing between embedded RNA molecules, transcript isoforms, including splice and length variants, as well as between alternative polycistronic transcripts. It has been demonstrated in several studies that long-read sequencing is able to circumvent these problems.

**Results:** In this work, we report the analysis of VZV lytic transcriptome using the Oxford Nanopore Technologies sequencing platform. These investigations have led to the identification of 114 novel transcripts, including mRNAs, non-coding RNAs, polycistronic RNAs and complex transcripts, as well as 10 novel spliced transcripts and 27 novel transcription start site isoforms and transcription end site isoforms. A novel class of transcripts, the nroRNAs are described in this study. These transcripts are encoded by the genomic region located in close vicinity to the viral replication origin. We also show that the VZV latency transcript (VLT) exhibits a more complex structural variation than formerly believed. Additionally, we have detected RNA editing in a novel non-coding RNA molecule.

**Conclusions:** Our investigations disclosed a composite transcriptomic architecture of VZV, including the discovery of novel RNA molecules and transcript isoforms, as well as a complex meshwork of transcriptional read-throughs and overlaps. The results represent a substantial advance in the annotation VZV transcriptome and in understanding the molecular biology of the herpesviruses in general.

## Background

Varicella zoster virus (VZV) is one of the nine herpesviruses that infect humans. It is the etiological agent of chickenpox (varicella) caused by primary infection and shingles (zoster) is due to reactivation from latency, which is established in the spinal and trigeminal ganglia [1]. VZV belongs to the *Varicellovirus* genus of the subfamily *Alphaherpesvirinae*, and is closely related to the pseudorabies virus (PRV) a veterinary *Varicellovirus* and to the herpes simplex virus type-1 (HSV-1) belonging to the *Simplexvirus* genus of alphaherpesviruses. The investigation of the VZV pathomechanism is not easy due to its highly cell-associated nature and the lack of animal models, except the SCID mouse model [2, 3]. The VZV virion is ∼80-120 nm in diameter and is composed of an icosahedral nucleocapsid surrounded by a tegument layer [4]. The outer virion component is an envelope derived from the host cell membrane with incorporated viral glycoproteins [5]. The VZV genome is composed of a ∼125-kbp double-stranded DNA molecule containing at least 70 annotated open reading frames (ORFs). The viral DNA consists of two unique genomic regions (UL and US), and a pair of inverted repeats (IRs) surrounding the US region. Three genes (ORF62, ORF63 and ORF64) located at the IR regions are therefore in duplicate [6, 7]. VZV encodes six genes (ORFs S/L, 1, 2, 13, 32, and 57), which are not present in HSV [8]. It has been long held that VZV does not express the RNA molecule homologues to the HSV latency-associated transcripts (LAT) [9], however, a recent study reported the identification of VZV latency-associated transcript (VLT) overlapping the ORF61 gene. It has also been thought that VZV lacks the α-TIF protein encoded by the vp16 gene that activates the transcription of immediate-early (IE) genes during the initial events of the virus life cycle [10]. VZV ORF10 together with ORF4, 62 and 63 are transcriptional transactivators that are all present in the virus tegument [11, 12]. ORF10 is supposed to substitute the function of vp16 in the transactivation of ORF62 (homologue of HSV ICP4), but unlike VP16, it is unable to control the expression of ORF4, ORF63, and ORF61 [13, 14].

The viral replication is primarily controlled by the regulation of transcription, which is carried out through a sequential activation of viral genes. First, the IE genes are expressed, which is followed by the activation of the early (E) and then the late (L) kinetic classes of genes [15]. The IE protein ORF62 is the major transactivator of the viral genes, which recruits the general transcriptional apparatus of the host cell and thereby controls the expression of other viral genes [16, 17]. The VZV E genes encode the proteins required for the DNA replication, while L genes code for the structural elements of the virus. Polycistronism represents an inbuilt characteristic of the herpesvirus genome [18,19]. However, the herpesvirus multigenic transcripts differ from the bacterial polycistronic transcripts in that the downstream genes on the viral RNA molecules are untranslated. An exception to this rule was found in the Kaposi’s sarcoma-associated herpesvirus [19, 20]. The functional significance of this organization of the herpesviral transcripts has not yet been ascertained. Alternative splicing expands the RNA and protein repertoire with respect of the one gene, one RNA/protein situation. In contrast to the beta and gammaherpesviruses, splicing events are rare among the alphaherpesviruses [21, 22]. In VZV only a few genes have been shown to produce spliced mRNAs so far, which include the ORF0 (also referred as ORFS/L) located at the termini of UL region, the UL15 homologue ORF42/45, the glycoprotein M (gM) encoding ORF50, and the newly discovered, multiple spliced VLT [6,9,23,74,75].

RNA editing includes the adenosine deaminase acting on RNA1 (ADAR1) enzyme that catalyzes the C-6 deamination of adenine (A) to inosine (I). The I is recognized as guanine (G) by the reverse transcriptase (RT) enzyme [26], therefore it can be detected with cDNA sequencing methods. In hyper-edited sites the ADAR1 transforms multiple adjacent As on an RNA strand. A G-enriched neighborhood has been described at the RNA editing sites [27] with an upstream uracil (U) exerting a stabilizing effect on the RNA-ADAR complex [28]. Additionally, RNA hyper-editing has been shown to play a crucial role in the viral life cycle by dodging the host’s immune system [29] and also in the control of DNA replication [30].

The translation of eukaryotic mRNAs occurs according to the scanning model, where the 40S ribosomal subunits scans the RNA strand in 5’ to 3’ direction and initiates the translation at the first AUG they encounter [31]. Mammalian mRNAs contain an essential sequence context for translation initiation, known as the Kozak consensus sequence [32]. A purine at −3 and G at +4 position has the strongest binding effect for the translational machinery, while AUGs with a different context tend to be overlooked by the ribosomal subunits (leaky scanning) [33]. Upstream ORFs in the 5’-untranslated regions (UTRs) of the RNAs have been shown to exert a regulatory effect on the protein synthesis through a process called translation reinitiation [31, 34]. In short upstream (u)ORFs translation reinitiation shows a positive correlation with distance between stop codon of uORF and the AUG of downstream ORF [35].

The hybridization-based microarray and the second-generation short-read sequencing (SRS) techniques have revolutionized genome and transcriptome research, including herpesviruses [36–39]. However, both techniques have limitations compared to long-read sequencing (LRS), including Pacific Biosciences (PacBio) and Oxford Nanopore Technologies (ONT) platforms. Microarray and SRS techniques perform poorly in the detection of multiple-intron transcripts, transcript length variants, and polycistronic RNA molecules, as well as transcriptional overlaps. Additionally, the LRS sequencing platforms are capable of determining the transcript ends with base-pair precision, without the need of any supplementary method.

Similarly to other LRS techniques, ONT cDNA sequencing is afflicted by sample degradation, resulting in false transcriptional start sites (TSSs). This problem can be mitigated by using cap selection. Another source of non-specificity is the presence of nucleotide sequences complementary to the MinION strand switching oligonucleotide, which can lead to either template switching [40, 41] or false priming [42]. False priming with the oligo(dT) primer can also occur at A-rich regions of the transcripts. These errors contribute to artifactual 3’-ends of the reads.

LRS techniques have already been used in genome and transcriptome studies of viruses [43–46], including herpesviruses [18, 21, 22, 47–49]. These reports have revealed a hidden transcriptome complexity, which especially included the discovery of long RNA molecules and a large variation of transcript isoforms. ONT and PacBio-based studies have also detected a number of embedded transcripts with in-frame truncated (t)ORFs in several herpesviruses, such as PRV [18, 43], HSV-1 [50], and human cytomegalovirus (HCMV) [21]. Additionally, a large extent of transcriptional read-through and overlaps has also been uncovered by these studies [18, 21, 22]. It has been proposed that the transcriptional overlaps may be the result of a transcriptional interference mechanism playing a role in genome-wide regulation of gene expressions [51].

In this work, we used the ONT MinION sequencing technique to investigate the structural aspects of the polyadenylated VZV transcripts. We report numerous novel transcripts, transcript isoforms, and yet unknown splice events. This study also explores a complex meshwork of transcriptional overlaps. Additionally, we report and analyze a hyper-editing event in a VZV transcript.

## Methods

### Cells and viral infection

Human primary embryonic lung fibroblast cell line (MRC5) was obtained from the American Type Culture Collection (ATCC) and grown in DMEM supplemented with antibiotic/antimycotic solution and 10% fetal bovine serum (FBS) at 37°C in a 5% CO_2_ atmosphere. The live attenuated OKA/Merck strain of varicella-zoster virus (VZV) was cultured at 37°C in MRC5 cell line, the cells were harvested by trypsinization, when the monolayers had displayed specific cytopathic changes. For subsequent propagation of the virus, infected cells were used to inoculate MRC5 cultures previously grown to full confluence at a ratio of 1:10 infected to uninfected cells. The cultures were then incubated at 37 °C for 5 days, when the cytopathic effect was near 100%.

### RNA purification

Total RNA was isolated immediately after collecting the infected cells, using the Nucleospin RNA Kit (Macherey-Nagel) according to the manufacturer’s instruction. Briefly, infected cells were collected by centrifugation and the cell membrane was disrupted using the lysis buffer provided in the kit. Genomic DNA was digested with the RNase-free rDNase solution (supplied with the kit). Samples were eluted in a total volume of 50μl nuclease-free water. To eliminate residual DNA contamination a subsequent treatment with the TURBO DNA-free Kit (Thermo Fisher Scientific) was conducted. The RNA concentration was measured with a Qubit 2.0 Fluorometer, with the Qubit RNA BR Assay Kit (Thermo Fisher Scientific).

### Poly(A) selection

Thirty-five μg of total RNA was pipetted in separate DNA LoBind Eppendorf tubes (Merck). The poly(A)^+^ RNA fraction was isolated from the samples using the Oligotex mRNA Mini Kit (Qiagen). RNA samples were stored at −80°C until use.

### Oxford Nanopore MinION cDNA sequencing and barcoding

The cDNA library was prepared using the Ligation Sequencing kit (SQK-LSK108; Oxford Nanopore Technologies) according to the modified 1D strand switching cDNA by ligation protocol. In short, the polyA(+) RNA fraction was reverse transcribed using an oligo(d)T-containing primer [(VN)T20 (Bio Basic, Canada). The RT reaction was carried out as recommended by the 1D protocol, using SuperScript IV enzyme (Life Technologies); a strand-switching oligo [containing three O-methyl-guanine RNA bases (PCR_Sw_mod_3G; Bio Basic, Canada)] was added to the sample. The cDNA was amplified by using KAPA HiFi DNA Polymerase (Kapa Biosystems) and Ligation Sequencing Kit Primer Mix (part of the 1D Kit) following the ONT 1D Kit’s manual. End repair was carried out on the samples using NEBNext End repair / dA-tailing Module (New England Biolabs), which was followed by “barcoding”: the C11 barcode (ONT PCR Barcoding Kit 96; EXP-PBC096) was ligated to the sample according to the 1D PCR barcoding (96) genomic DNA (SQK-LSK108) protocol, Barcode Adapter ligation step. The barcoded-sample was amplified by PCR using KAPA HiFi DNA Polymerase, as well as the C11 PCR barcode according to the 1D PCR barcoding protocol. The PCR product was end-repaired, then it was followed by adapter ligation using the sequencing adapters supplied in the kit and NEB Blunt/TA Ligase Master Mix (New England Biolabs). The cDNA sample was purified between each step using Agencourt AMPure XP magnetic beads (Beckman Coulter). The concentration of cDNA library was determined using a Qubit 2.0 Fluorometer through use of the Qubit (ds)DNA HS Assay Kit (Thermo Fisher Scientific). Samples were loaded on R9.4 SpotON Flow Cells, while the base calling was performed using Albacore v2.1.10.

### Cap selection, cDNA synthesis and sequencing

For the precise determination of TSSs, the ONT’s 1D strand switching cDNA by ligation protocol was combined with a 5’-cap specific protocol. The cDNA sample was prepared from total RNA using the TeloPrime Full-Length cDNA Amplification Kit (Lexogen). Reverse transcription (RT) reaction was performed at 46°C for 50min according to the kit’s recommendations by using an oligo(d)T-containing primer (5’ −> 3’: TCTCAGGCGTTTTTTTTTTTTTTTTTT). The RT product was purified by using spin column-based method (silica columns are from the Lexogen kit). A double-strand specific ligase enzyme (Lexogen Kit) was used to ligate an adapter to the 5’C of the cap of the RNA. The reaction was carried out at 25°C, overnight. The sample was washed by applying the silica-column method. The Second-Strand Mix and the Enzyme Mix (both from the TeloPrime Kit) as well as a Verity Thermal Cycler (Applied Biosystems) were used to produce double-stranded cDNAs according to the Kit’s guide. In order to generate sufficient amount of cDNAs for MinION library preparation, samples were amplified by endpoint PCR following the TeloPrime Kit’s manual. The generation of the sequencing-ready library from this sample is based on the 1D strand switching cDNA by ligation protocol from ONT. The Ligation Sequencing kit (SQK-LSK108, ONT) and NEBNext End repair / dA-tailing Module (New England Biolabs) was used to repair the cDNA ends. This step was followed by the 1D adapter ligation, which was carried out according to the 1D protocol, using the NEB Blunt/TA Ligase Master Mix (New England Biolabs). We used barcoding for the better *in silico* identification of the transcripts’ ends. We found this method useful because it helped the base-pair precision mapping of TSSs. The library was sequenced on an ONT R9.4 SpotON Flow Cells.

### Data analysis and alignment

Reads resulting from ONT sequencing were aligned to the reference genome of VZV (GeneBank accession: NC_001348.1) and the host cell genome (Homo sapiens - GRCh37, BioProject number: PRJNA31257) using GMAP v2017-04-24[53]. In order to annotate the 5’ ends, the last 16nt of the MinION 5’ strand switching adapter or the last 16nt of the cap selection adapter was aligned in a window of −10 nt and +30 nt from the first mapped position of a read using the Smith-Waterman algorithm, with a match cost of +2, a mismatch cost of −3, a gap opening cost of −3, and a gap extension cost of −2. Read ends with a score below 14 were considered false 5’ ends and were discarded. A 5’-end position was considered a TSS if the number of reads starting at this position was significantly higher than at other nucleotides in the region surrounding this start position. For this the Poisson-probability (Poisson[k_0_;λ]) of k_0_ read starting at a given nucleotide in the −50 nt to +50 nt window from each local maximum was calculated with 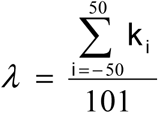. The 5’-ends of the low-abundant long reads were inspected individually using the Integrative Genome Viewer (IGV) [54]. Poly(A) tails were defined using the same algorithm and parameters, by mapping 15 nt of homopolymer As or Ts to the soft-clipped region on the read’s end. Read ends with a score below 14 or with more than 5 As/Ts directly upstream their position were considered artefacts of false priming, and were discarded. Transcription end sites (TESs) were defined using the same criteria for the Poisson-probability as in case of TSSs. Reads passing both criteria and present in more than three copies were considered as “certain transcript isoforms”. Those with less than three copies or the longest unique reads with questionable 5’ ends were considered “uncertain transcript isoforms”. Reads with sequencing adapters or poly(A) tails on their both ends were discarded, except for complex transcripts, which were individually inspected using IGV. Reads with a larger than 10nt difference in their 5’ or 3’ ends were considered novel length isoforms (L: longer 5’ UTR, S: shorter 5’ UTR, AT: alternative 3’ termination). Short length isoforms harboring a truncated version of the known open reading frame (ORF) were considered novel putative protein coding transcripts, and designated as ‘.5’. If multiple putative protein coding transcripts were present, then the one with the longest 5’ UTR was designated ‘.5’ and its shorter versions were labelled in an ascending order. Transcripts with TESs located within the ORFs of genes (therefore lacking STOP codons) or with TSSs within the coding regions without in-frame ORFs were both considered non-coding transcripts.

Multigenic transcripts containing at least two genes standing in opposite were named complex transcripts. If a TSS was not obvious at these transcripts, we assumed that they start at the closest upstream annotated TSSs. Splice junctions were accepted if the intron boundary consensus sequences (GT and AG) were present in at least ten sequencing reads and if the frequency of introns was more than 1% at the given region.

To assess the homology of protein products possibly translated from the ORFs of novel transcript isoforms we used the online BLASTP suit[55], with an expected threshold of 10.

To evaluate the effect of hyper-editing on the secondary structure of RNAs, we used the RNAstructure Software suite [56] with the following parameters: temperature=310,15 K, Max % Energy Difference=10, Max Number of Structures=20, Window Size=0. To simulate the presence of inosine, we changed the edited adenines to guanine. To assess the secondary structure of the ORF62-AS2-ORF62 dsRNA hybrid we used the first 467 bases of the ORF62 labeled ORF62-5’ (from genomic position 109,204 to 108,738) as one of our starting sequences, and the whole sequence of ORF62-AS2 as the other one. Reads were visualized using the Geneious [57] software suite and IGV. GC-, CCAAT- and TATA-boxes were annotated using the Smith-Waterman algorithm for canonical motifs [58].

## Results

### Analysis of the VZV transcriptome with third-generation sequencing

In this work, the ONT MinION RNA-Seq technique was used for profiling the genome-wide expression of the lytic VZV transcripts (**Figure 1**). Both cap-selected and non-cap-selected protocols were applied for the analyses. Sequencing of the non-cap-selected sample yielded 511,886 reads of which 57,888 mapped to the VZV reference genome (GeneBank Accession: NC_001348.1) with an average mapped read length of 1,349 bp, and an average coverage of 625, while 453,998 reads mapped to the host cell genome (Homo sapiens - GRCh37, BioProject number: PRJNA31257). Sequencing of the cap-selected samples yielded 827,608 reads, of which 509,531 mapped to the viral genome, with an average coverage of 1,200, and 318,077 reads mapping to the host cell. This latter technique performed poorly in VZV, which was indicated by the short average aligned read length (294 bp) (**Figure 2 panel a.**). Intriguingly, we obtained the same poor result with PRV [22] and HSV-1 (our unpublished results), however, the cap-selection technique performed very well in the analysis of the transcriptomes of HCMV, the baculovirus *Autographa californica* multiple nucleopolyhedrosis virus (AcMNPV) and vaccinia virus (our unpublished results). These results together with an analysis of the nucleotide distribution at the vicinity of the 5’-ends in the cap-selected datasets (**Figure 2 panel b.**) demonstrate that the read length of cap-selected RNA-s is not determined by the GC-content (PRV: 73%, HSV-1: 68%, VZV: 46%), but by a yet unknown factor, which is common in alphaherpesviruses.

**Figure 1.**
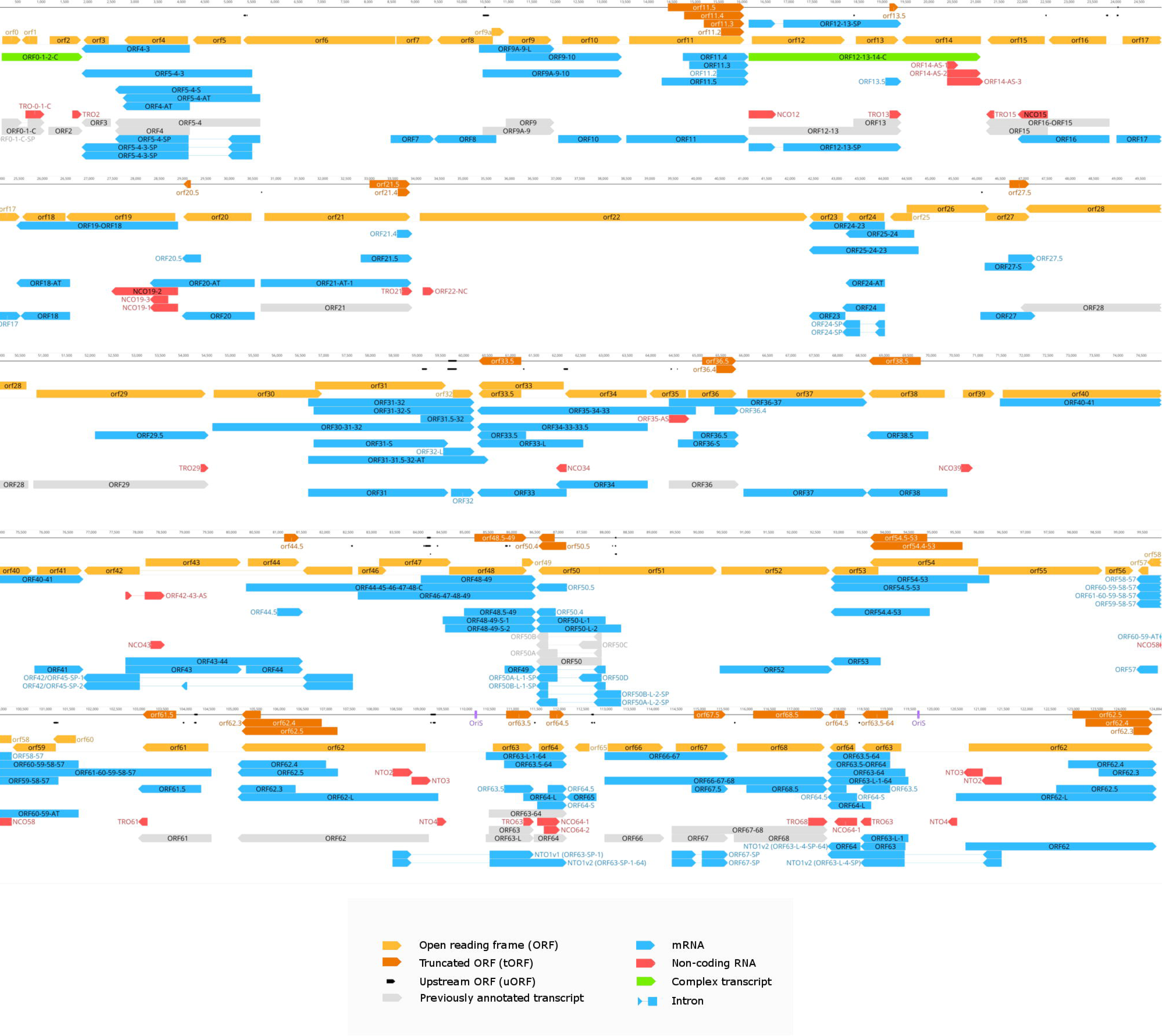
The VZV transcriptome. Color code: light orange arrow-rectangles: open reading frames (ORFs); black arrow-rectangles: upstream ORFs; dark orange arrow-rectangles: truncated (t)ORFs; gray arrow-rectangles: previously annotated transcripts; blue arrow-rectangles: novel mRNAs; red arrow-rectangles: novel ncRNAs; green arrow-rectangles: novel complex transcripts.

**Figure 2.**
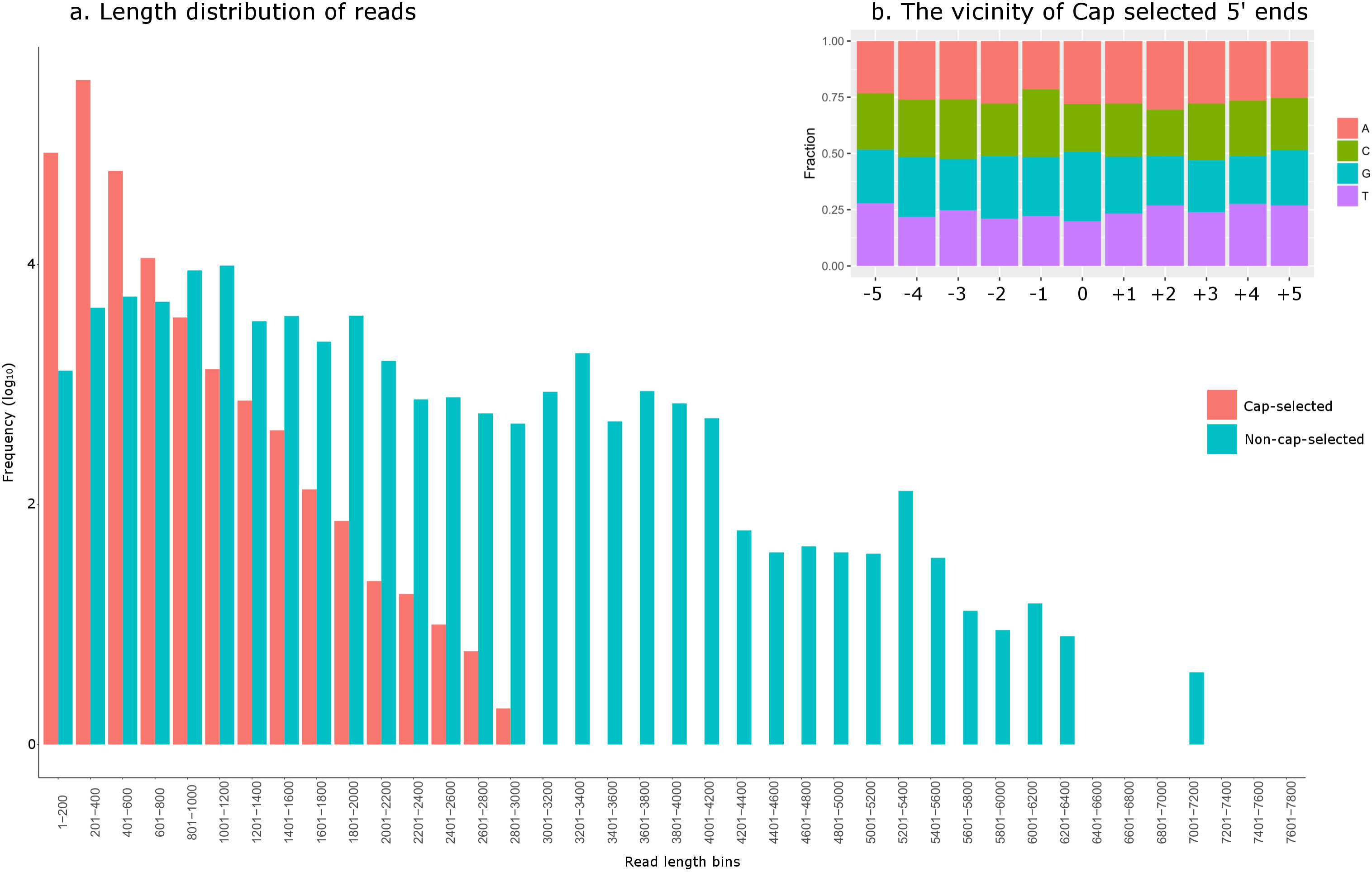
The read length distributions. a. The frequency of the reads of the cap-selected and non-cap-selected sequencing binned by 200 nucleotides shows that the cap-selected reads are skewed towards shorter read lengths. b. The frequency of each nucleotide in the vicinity of the 5’ ends of the cap-selected reads shows random distribution, suggesting that the short read lengths are not caused by the presence of a specific nucleotide on the RNA but other, yet unknown reason.

In total, we detected 1,124 5’-ends and 255 3’-ends in the non-cap-selected dataset, and 1,472 5’-ends and 279 3’-ends in the cap-selected dataset. The number of 5’-ends qualifying as a TSS was 10.86% of the total 5’-end positions, while 32.95% of the total 3’-end positions turned out as TES (Table 1). We excluded 49 5’-ends and 16 3’-ends from the non-cap-selected and 22 5’-ends and 16 3’-ends from the cap-selected datasets because they proved to be the products of false priming.

**Table 1.**
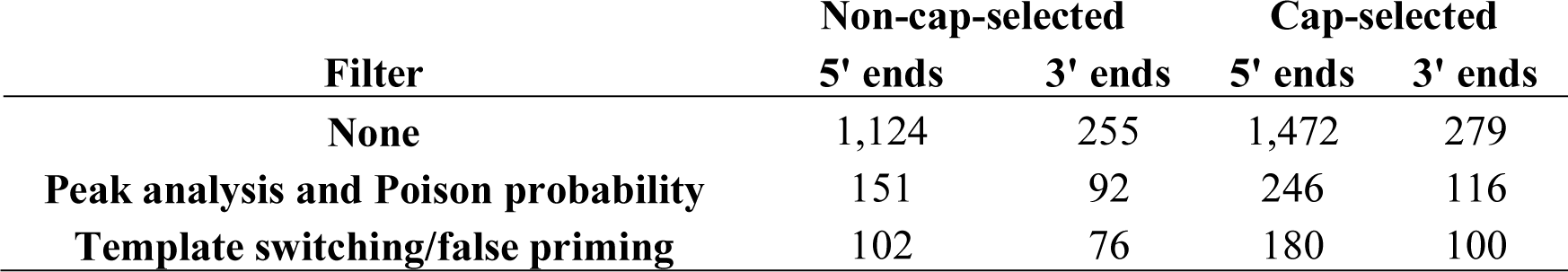
The number of read ends following each step of filtering.

Earlier results with primer extension, S1 nuclease assay and Illumina sequencing determined the transcript ends of some of the RNA molecules of VZV. Using the ONT sequencing platform, we confirmed 18 previously known TSSs and nine TESs. Additionally, we annotated 124 new TSSs and 71 new TESs listed in **Supplementary Table 1**. The sequences upstream of the TSSs and TESs were analyzed *in silico* to detect putative GC-, CCAAT- and TATA-box motifs, and Poly(A) signals (PASs), by motif searching (**Table 2**).

**Table 2.**
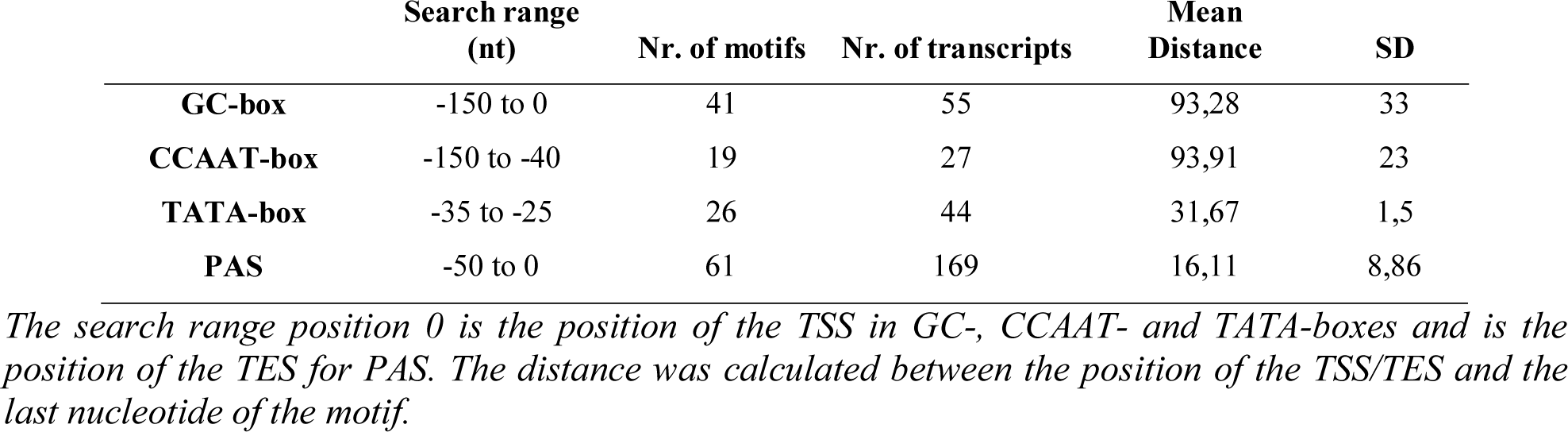
Putative promoter sequences and Poly(A) signals (PASs) of the VZV transcripts.

In this work, we detected an enrichment of As and Us in an interval of up to −10 nt upstream of the TES, and an abundant GT-rich region downstream in the +10 nt interval (**Figure 3.**). We annotated altogether 246 novel transcripts of which 149 were confirmed by at least three reads, listed in **Supplementary Table 2** and **Supplementary Table 1**. Additionally, we mapped the TSS and TES of 37 previously detected transcripts lacking former mapping of RNA end positions (**Supplementary Table 1**).

**Figure 3.**
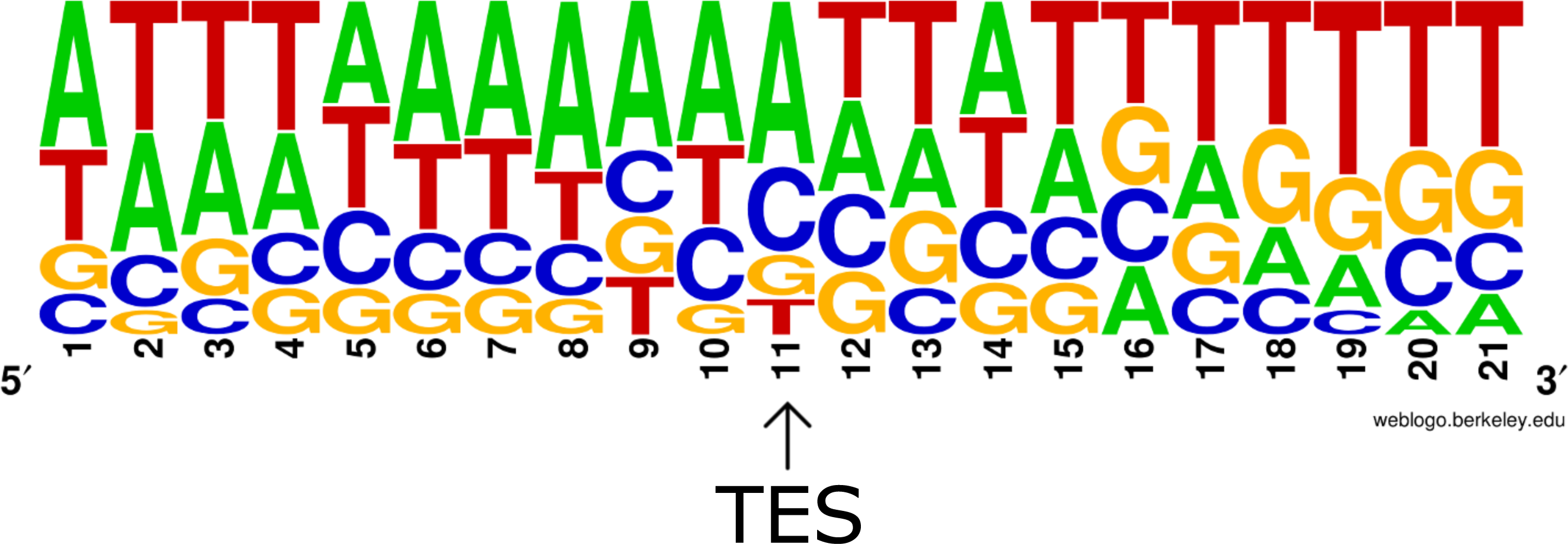
The frequency of nucleotides in the vicinity of transcriptional end sites (TES). The weblogo shows an enrichment of A and U bases upstream, while an enrichment of G and U bases downstream of the TES. This pattern is akin with the sequence surroundings of mammalian TESs.

### Putative mRNAs

Transcripts embedded into a larger RNA molecules are easily detected by the LRS techniques. If such a transcript contains an in-frame ORF shorter [truncated (t)ORF] than that if the host ORF, it can be considered as a putative mRNA potentially coding for an N-terminally abridged polypeptide. Earlier *in silico* approaches have annotated the VZV ORFs [6, 7] one of which (the orf33.5) is a tORF. In this study, we report the identification of 28 embedded transcripts containing tORFs. We could identify promoters for 12 of these transcripts, located at a mean distance of 93.85±32.18 nt (mean ± SD) from their TSSs (**Supplementary Table 3**). Three of these transcripts (ORF13.5, ORF54.5-53 and ORF62.3) contain strong Kozak consensus sequences near their AUGs, while 19 have at least one of the two essential nucleotides present on their −3 or +4 positions. Twenty-five of the possibly protein coding transcripts contain a PAS at a 19.44±8.91 (mean ± SD) distance upstream their TESs (**Supplementary Table 3**).

**Table 3.**
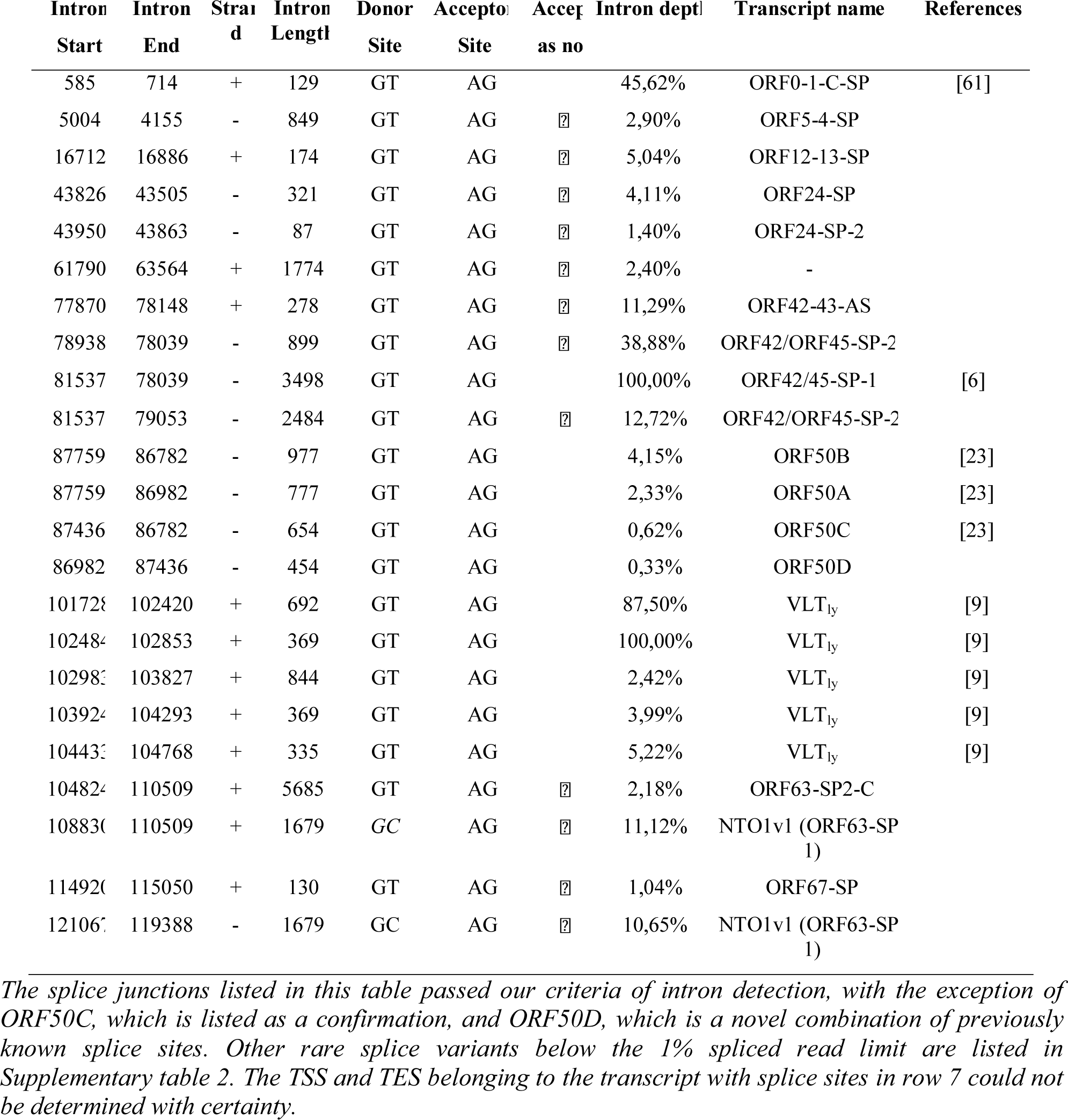
Splice junction sites of the VZV transcriptome.

### Determination of the ends of mRNAs with annotated ORFs

In this work, we determined the TSSs and TESs of 25 mRNAs whose ORFs had earlier been described (**Figure 1 Supplementary Table 1).** The TSS and TES of these monocistronic mRNAs were not characterized before by any other means. Twelve of these transcripts have a promoter sequence 51.57±33.31 (mean ± SD) upstream their TSSs, and 19 have a canonical PAS upstream their TESs at a mean distance of 14.96±11.59 (mean ± SD).

### Non-coding transcripts

The ncRNAs are transcripts without the ability to encode proteins. They can overlap the coding sequences of the genes in the same orientation, or in the opposite orientation [antisense (as)RNAs], or they can be located at the intergenic regions. The 5’-truncated (TR) transcripts have their own promoters but share the PASs and TESs with the ‘host’ transcript, while the 3’-truncated (NC) RNAs are controlled by the same promoters as the canonical transcript in which they are embedded but have no in-frame ORFs. Twenty-three of the novel non-coding transcripts are long non-coding (lnc)RNAs exceeding 200 bps in length per definition, while five of them are small non-coding (snc)RNAs with a size below 200 bps.

In this work, we identified 20 embedded ncRNAs, of which 8 were 5’-truncated, while 12 were 3’-truncated transcripts (**Supplementary Table 1**). We also detected one intergenic and seven antisense ncRNAs, all of them being controlled by their own promoters. In total, 17 of the novel ncRNAs contain a canonical promoter sequence 60.4±35.19 nt (mean ± SD) upstream their TSSs, while 21 have a PAS at a 16.35±8.87 (mean ± SD) distance of their TESs. ORF42-43-AS and ORF35-AS overlap multiple mRNAs. ORF42-43-AS stands in antisense orientation with respect of ORF42/ORF45-SP-1 and ORF42/ORF45-SP-2, while in tail-to head orientation with ORF43 and ORF43-44. ORF35-AS stands in antisense orientation with the polycistronic transcript ORF35-34-33 and in tail-to-head orientation with ORF36 and ORF36-S and ORF36-37. (**Figure 1**). The LAT RNA has been described in every member of the alphaherpesvirus subfamily [59, 60], including VZV [9]. We confirmed the existence of four previously detected lytic isoforms of VLT (VLT_ly_) of which two are TSS isoforms, one is a splice variant and one is both a TSS isoform and a splice variant (**Figure 4, Supplementary Table 2**).

**Figure 4.**
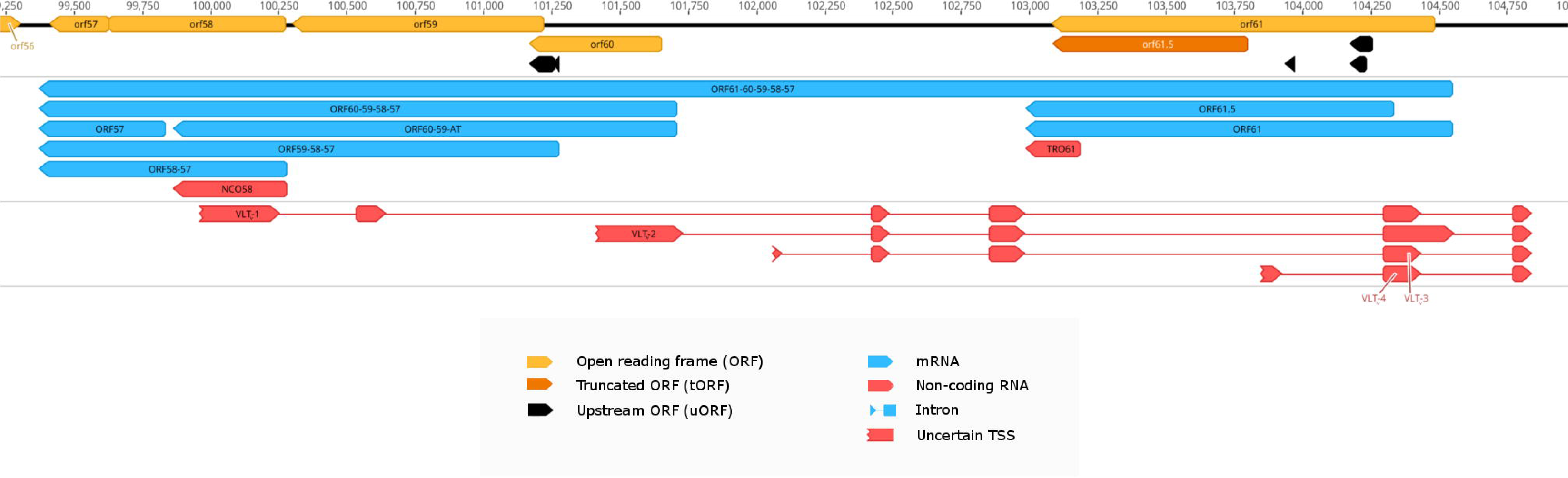
The genomic region of VLT. Color code: light orange arrow-rectangles: open reading frames (ORFs); dark orange arrow-rectangles: truncated (t)ORFs; black arrow-rectangles: upstream ORFs; gray arrow-rectangles: previously annotated transcripts; blue arrow-rectangles: novel mRNAs; red arrow-rectangles: ncRNAs; Reads of the VLT_ly_ transcript isoforms were present in very low abundance, thus their 5’ ends are feathered resembling uncertain TSSs.

### 5’- and 3’-UTR isoforms

The 5’-UTR isoforms (TSS variants) start upstream or downstream of the TSS of the earlier annotated transcripts, and their expression is regulated by their own promoters, while 3’-UTR isoforms (TES variants) contain distinct PAS and their polyadenylation occurs upstream or downstream of the TES of the main transcript. In this report, we detected 19 novel 5’-UTR length variants, 7 being shorter and 12 being longer than the earlier annotated transcript isoforms. We found canonical promoter sequences in 10 of the 5’-UTR isoforms at a distance of 94.93±38.58 (mean ± SD). Additionally, we detected 8 3’-UTR length isoform, all with a canonical PAS 22±7.65 (mean ± SD) upstream their TESs. An intriguing finding is the putative TSS at the genome position +4 belonging to the rare abundance transcript ORF0-1-L-C and ORF0-1-2-L-C at the extreme termini of UL region (TR_L_), which suggests the existence of a promoter located on the other terminal repeat (TR_S_) of the genome. This putative promoter is supposed to be active in the circular genome.

### Splice sites and splice isoforms

Reverse transcription can produce false introns between repetitive sequences of the template RNA due to the phenomenon of template switching. In order to exclude these artefacts, we removed sequencing reads with low abundance (≤ 1%) and with a repeat of more than six nucleotides next to one of the splice junctions. From the initial set of 10,064 unique splice acceptor and donor site candidates 24 matched our criteria resulting in a total of 16 splice isoforms being above the 1% intron depth level of acceptance. We detected two novel splice variants encoded by the ORF42/45 gene. Furthermore, we detected twelve novel splice sites and confirmed the existence of nine previously described spliced transcripts, all with a consensus GT at the splice donor site and AG at the splice acceptor site.

ORF63, homologue to the HSV-1 and the PRV us1 gene, is one of the main regulators of VZV transcription. In this work, we discovered ten novel splice variants of ORF63. Similarly to the HSV-1 US1 mRNA, NTO1v1 and the NTO1v3 harbors one intron in its 5’ UTR, while the NTO1v4 is spliced twice just as the PRV US1 mRNA [47, 50] (**Figure 5**). Ten splice junction of ORF63 splice variants coincide with those of the VLT_ly_ splice variants [9], thus they were labeled as VLT-ORF63-C (**Figure 6**). One of the splice donor sites, present in both NTO1v1 and NTO1v2 is GC, which is different from the canonical splice donor sequence. We detected all of the three ORF50 splice isoforms previously described, but ORF50C occurs in relatively low abundance, below our acceptance threshold. We detected another splice variant in low abundance in the ORF50 cluster, and named it ORF50D (Table 3, Figure 7).

**Figure 5.**
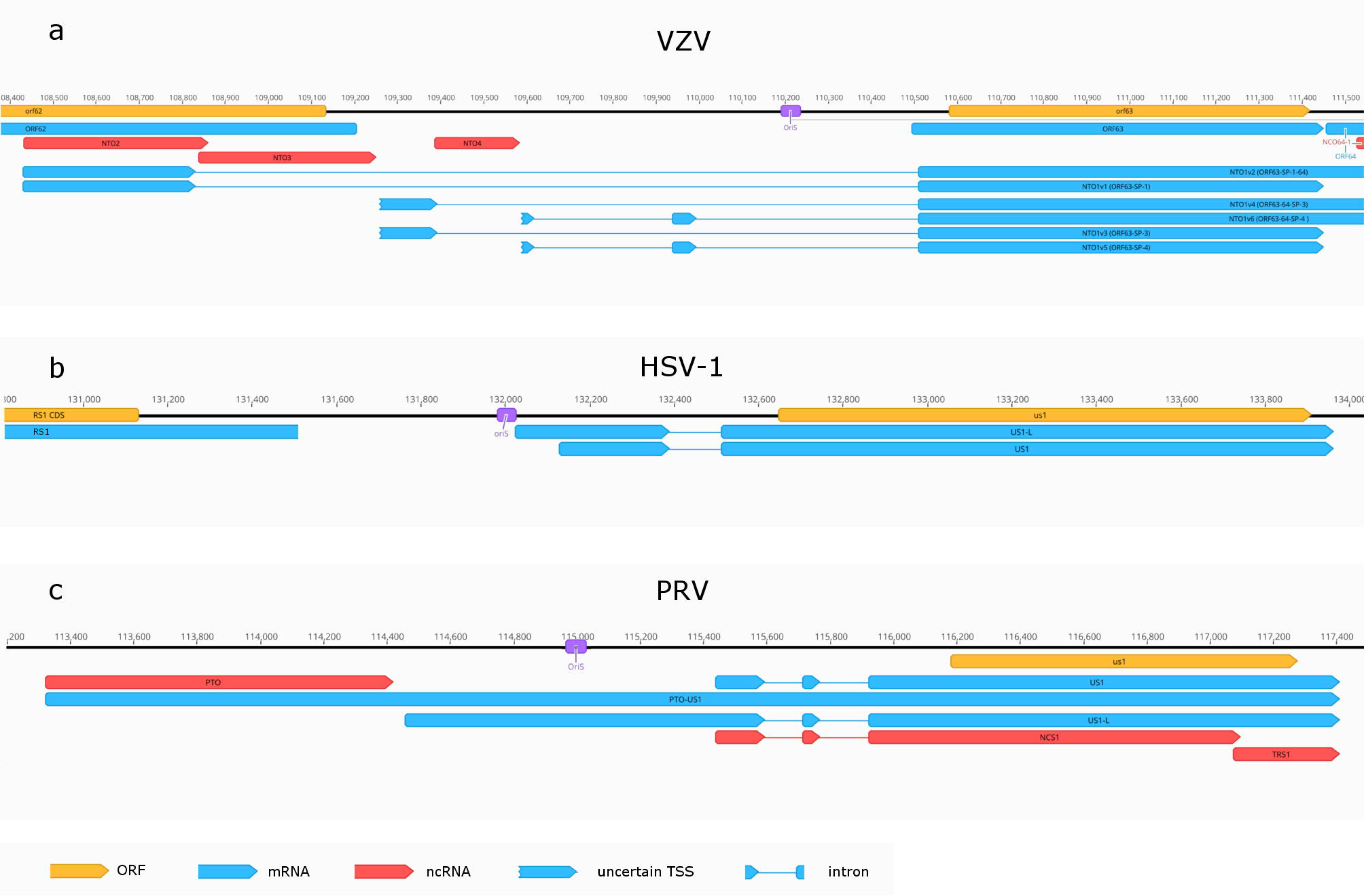
Structurally similar regions of three alphaherpesvirus transcripts neighboring OriS. Color code: light orange arrow-rectangles: open reading frames (ORFs); blue arrow-rectangles: mRNAs; red arrow-rectangles: ncRNAs. The ORF63 of VZV, similarly to the HSV-1 US1 has a two-exon-baring splice isoform, while akin with PRV’s US1 the three-exon-baring splice isoforms. Both VZV and PRV express non-overlapping non-coding RNAs in the proximity of OriS.

**Figure 6.**
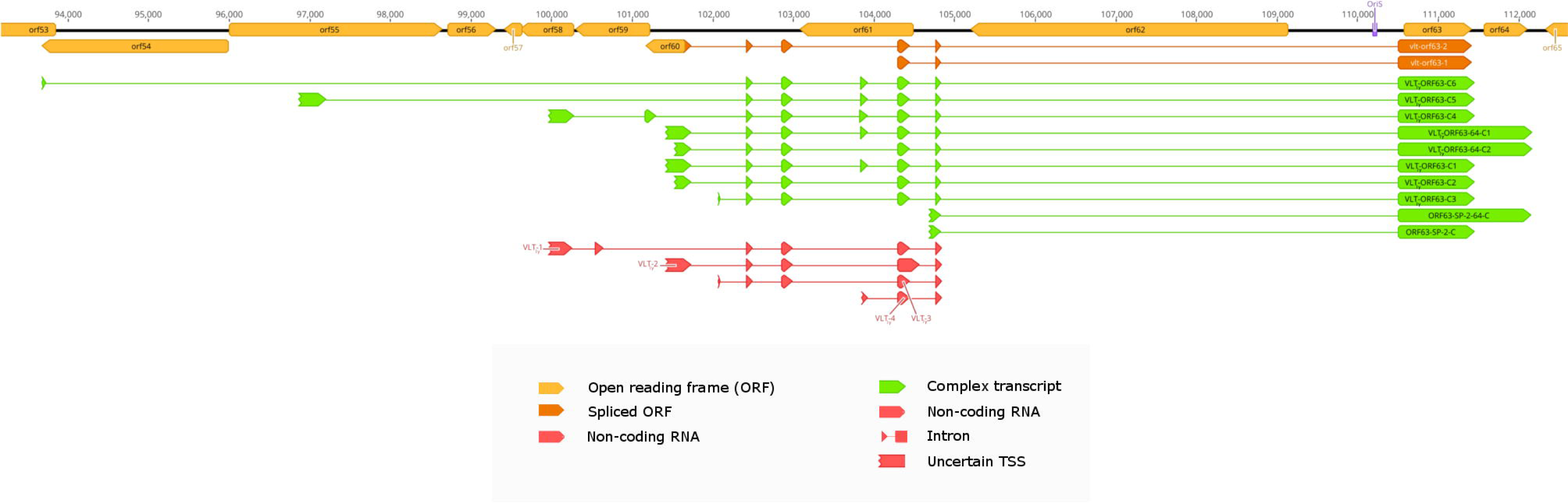
Splice isoforms of ORF63 and ORF64 with a similar splicing pattern as the VLT_ly._ Color code: light orange arrow-rectangles: open reading frames (ORFs); dark orange arrow-rectangles: spliced ORFs; green arrow-rectangles: complex transcripts; red arrow-rectangles: ncRNAs. AUGs present in the upstream exons of the ORF63 and ORF64 splice isoforms encompasses two spliced ORFs, the vlt-orf63-1 translated to a hypothetical protein product 88 aa longer, while the vlt-orf63-2 translated to a hypothetical protein product 179 aa longer than orf63.

**Figure 7.**
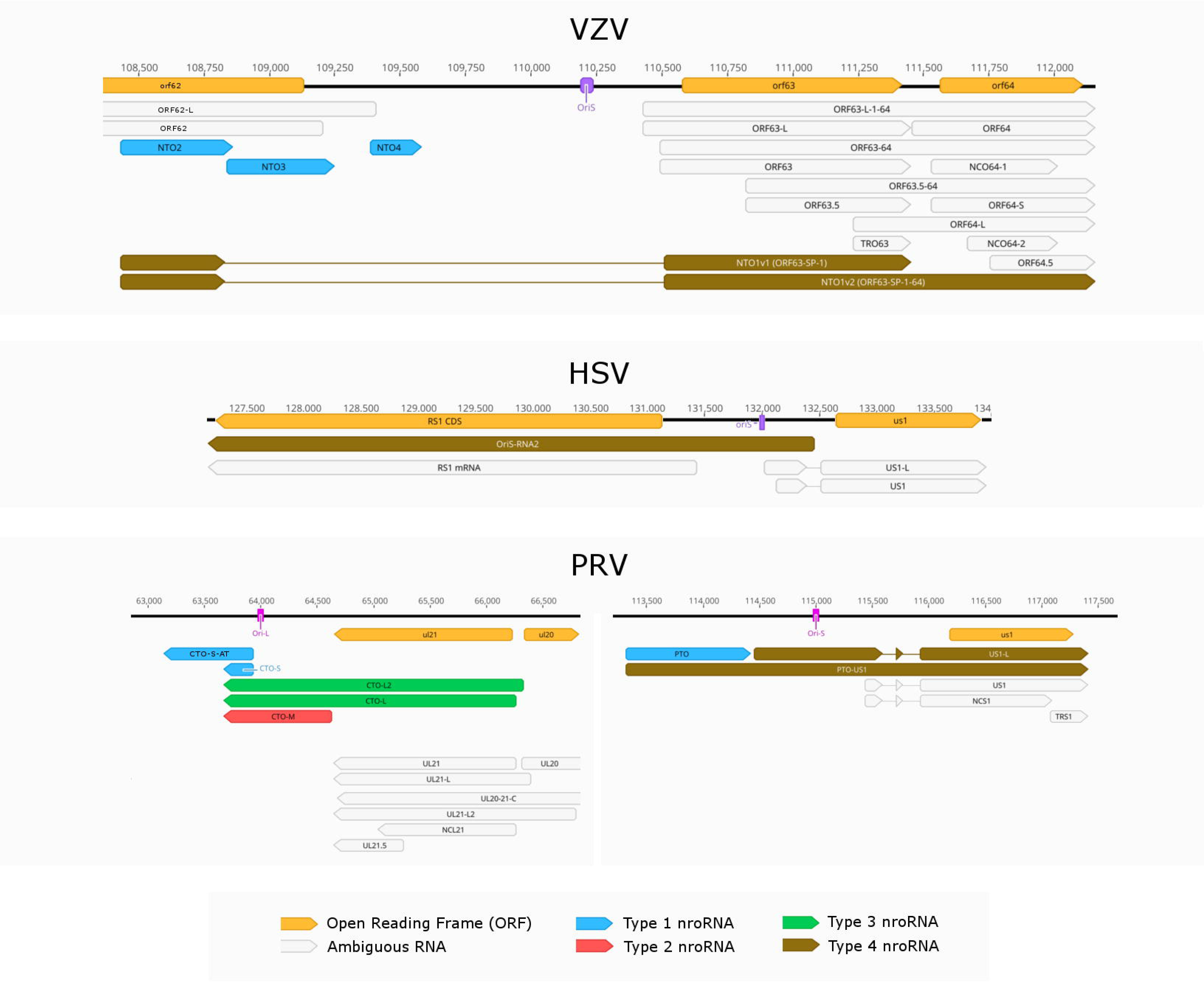
Types of near-replication-origin (nro)RNAs of three alphaherpesviruses. Color code: light orange arrow-rectangles: open reading frames (ORFs gray arrow-rectangles: ambiguous RNAs; blue arrow-rectangles: ncRNAs not overlapping the Ori (type 1); red arrow-rectangles: ncRNA overlapping the Ori (type 2); green arrow-rectangles: mRNAs overlapping the Ori and with long 3’ UTRs (type 3); brown arrow-rectangles: mRNAs overlapping the Ori and with long 5’ UTRs (type 4).

In four splice isoforms splicing effects the length of the ORFs, all producing N-terminally truncated hypothetical isoforms of previously detected proteins (Table 4).

**Table 4.**
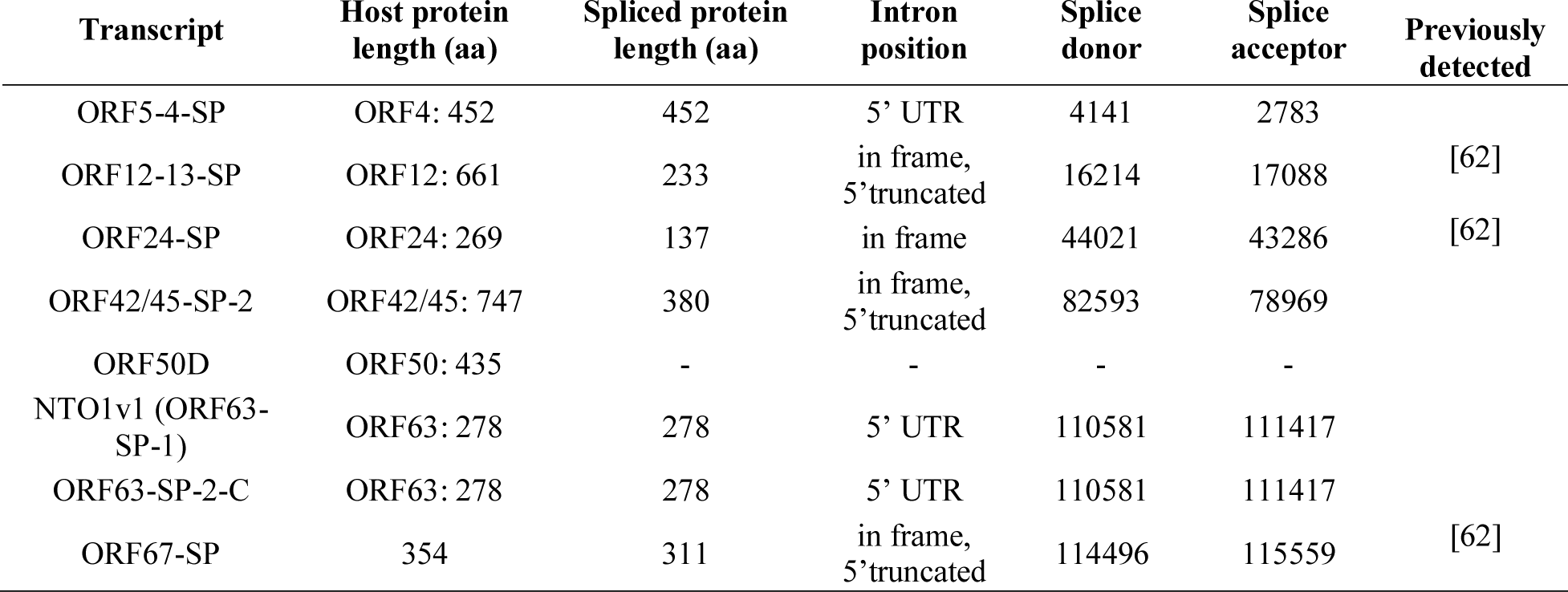
The proteins and hypothetical proteins produced by spliced transcripts.

### Near-replication-origin (nro)RNAs – a novel class of transcripts

In our earlier work, we reported [63] that PRV expresses transcripts located near the replication origins (Oris): the CTO family (including CTO-S, CTO-S-AT, CTO-M and CTO-L) at the OriL and the PTOs (PTO and PTO-US1) [18] at the OriS. Expression of nroRNAs has also been described in HSV-1 (OriS-RNS: [64]) and HCMV (OriLyt: [65]; RNA4.9: [66]). The VZV genome contains exclusively OriS, and lacks the replication origin at the UL segment. We identified 9 nroRNAs starting in the proximity of VZV OriS. The NTO1v1, NTO1v3 and NTO1v5 are the spliced long TSS variants of ORF63 while the NTO1v2, NTO1v4 and NTO1v6 are the spliced long TSS variant of ORF64. NTO2 starts at the same TSS as NTO1v1 but terminates 30 bp downstream of its splice donor site. The NTO3 and NTO4 are positioned downstream of NTO2. A similar transcriptional arrangement can be observed in PRV, where the TSS of the PTO-US1 (positionally similar to the NTO1v1) is located at the same genetic locus as PTO (positionally similar to the NTO2, NTO3 and NTO4, but not homologous) (**Figure 5 panels a and c**). Based on this result, we can distinguish four types of nroRNAs: (1) ncRNAs that do not overlap the Ori (such as CTO-S, CTO-S-AT, PTO, as well as NTO2, NTO3 and NTO4); (2) ncRNAs that do overlap the Ori (such as CTO-M), (3) mRNA isoforms with very long alternative TES (such as CTO-L and CTO-L2); and (4) mRNA isoforms with very long TSS variant [such as PTO-US1, US1-L (PRV), OriS-RNA2 (HSV-1), and the now discovered NTO1 isoforms] (**Figure 7**).

### Polycistronic and complex transcripts

A major issue of SRS, as wells as microarray and quantitative PCR approaches is that they have severe limitations in distinguishing between mono- and polycistronic transcripts. In contrast, LRS sequencing is suitable for making this distinction, and it is particularly superior in the detection of low abundance multi-genic transcripts. In this work, we identified 33 novel multigenic RNAs, including 29 polycistronic and 4 complex transcripts. Complex transcripts are multigenic RNAs that contain one or more genes in opposite orientations. Antisense sequences on the complex transcripts are unable to encode proteins. We also detected ten complex transcripts in low abundance in the region of VLT, seven of which are co-terminal with ORF63 and three with ORF64. These transcripts overlap with several oppositely oriented coding sequences and are spliced in a similar manner as the VLT_ly_ isoforms (**Figure 6, Supplementary Table 2**). *In silico* analysis detected an in-frame ORF incorporating the coding sequence of ORF63 (it has an upstream AUG). This results in VLT_ly_-ORF63-C1, VLT_ly_-ORF63-C4, VLT_ly_-ORF63-C5, VLT_ly_-ORF63-C6 and VLT_ly_-ORF63-64-C1 encoding hypothetical proteins whose N-terminal is longer with 88 amino acids (aa) than the one encoded by the orf63 gene, while the VLT_ly_- ORF63-C2; VLT_ly_-ORF63-C3 and the VLT_ly_-ORF63-64-C2 encoding hypothetical proteins with 179 aa longer than those coded by orf63 (**Figure 6**), The N-terminal overhangs of the putative proteins show no homology with any other known proteins in online databases.

### Transcriptional overlaps

Transcripts can overlap each other in parallel (tandem or head-to-tail), convergent (tail-to-tail) or divergent (head-to-head) manner (**Figure 8**). RNAs identified and annotated with a certain TSS and TES in this work form a total of 486 overlaps, of which 16 are head-to-head, 454 are head-to-tail and 16 are tail-to-tail (**Supplementary Table 4**). The overlaps can be full or partial. Full overlaps can be formed between the RNA molecules encoded by polycistronic transcription units, between embedded and host mRNA molecules, between mRNAs and 3’ as well as 5’ truncated transcripts, between the TSS and TES isoforms, etc. Partial overlaps can be formed between every transcript type. An overlap is ‘hard’ if two genes can only produce overlapping transcripts, or ‘soft’ if both overlapping and non-overlapping transcripts are formed. The soft overlap can be the result of alternative promoter usage (TSS isoforms) or transcriptional readthrough (TES isoforms). An example for the latter case is the ORF17 and ORF18, which produce non-overlapping transcripts, but the TES isoform of ORF18 (ORF18-AT), with its additional 89 bps in the 3’UTR, overlaps with ORF17 in a tail-to-tail manner (**Figure** ### panel c.).

**Figure 8.**
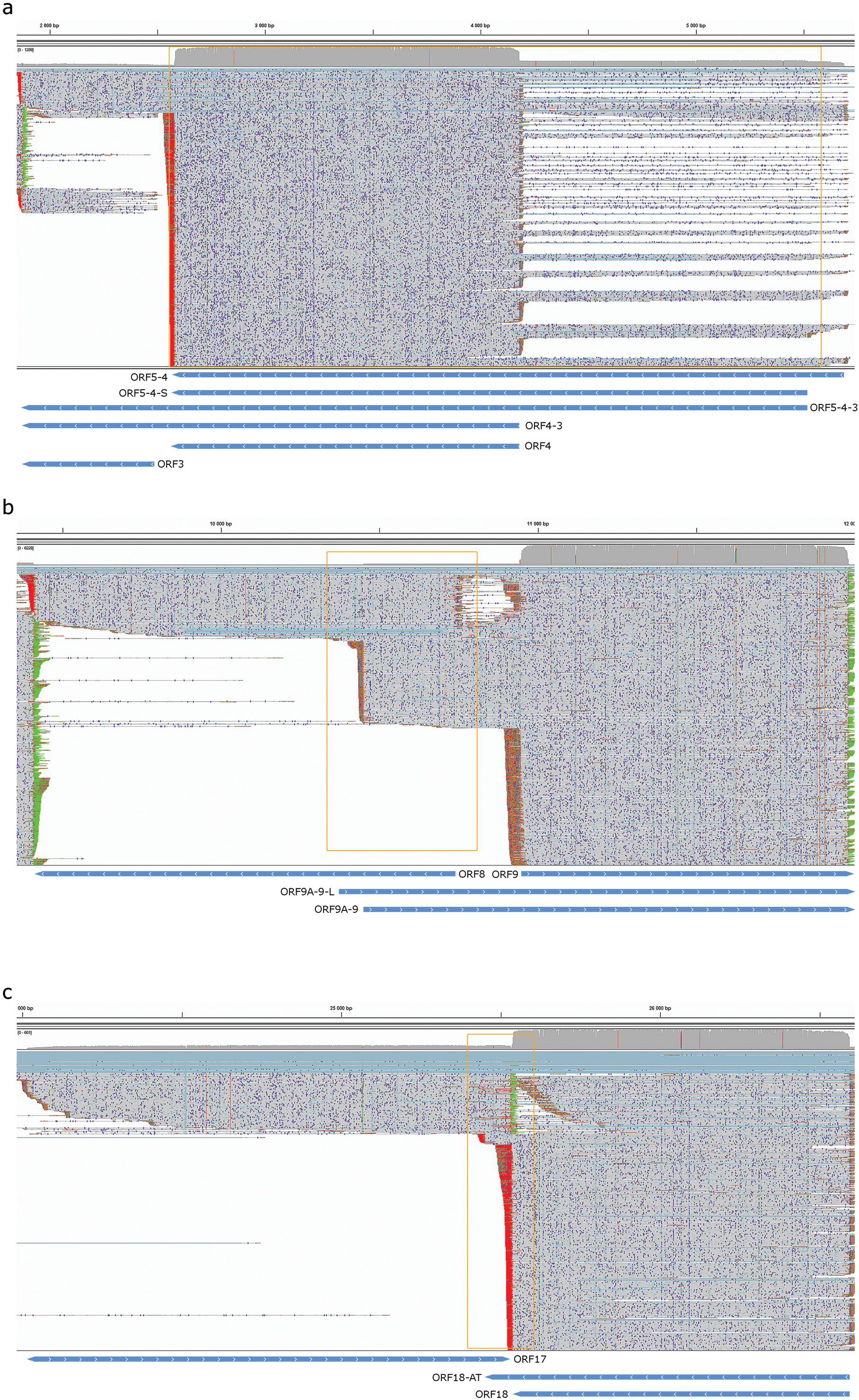
Types of overlaps between VZV transcripts. a. Parallel overlaps of the ORF5-4-3 cluster. b. Divergent overlaps of the ORF8 and ORF9-L and ORF9A-9 transcripts. c. Convergent overlaps of the ORF17 and ORF18-AT transcripts.

### Upstream ORFs

Using *in silico* methods, we detected 44 potential (u)ORFs on the 5’-UTRs of 81 VZV transcripts **(Figure 1)**. Five of the longer TSS variants contain uORFs, while their shorter isoform does not. We have previously described this phenomenon in HCMV [49] and PRV [22]. The average size of an uORF was 54.9 ± 44.64 nt (mean ± SD, median=35 nt), while the average distance of the uORF stop codon from the protein coding ORF’s AUG was 174.84 ± 143.58 nt (mean ± SD, median=110 nt). This space between the two uORF and the canonical ORF is enough for a potential reinitiation event. We identified a Kozak consensus sequences in five uORFs.

### RNA editing

The sequencing reads of NTO3 transcript show a very high frequency of A to G substitution, which is not present in the overlapping reads of other transcripts. We found that 58% of all substitutions are A->G (**Figure** 9 panel b) in the reads of NTO3, which is significantly higher than the 12.98% in the overlapping transcripts in the same region (p<0.0001, Fisher’s exact test) (**Figure 9 panel a and c**), making 22.07% of all As of the transcript edited. This suggest a hyper-editing event in NTO3. Using the MEME software suite [67] we could detect a slight enrichment of the GU dinucleotide preceding the editing site resulting in a GUAG motif (the editing site is underlined) (**Figure 8 panel d**). Using the RNAstructure software suite [56], we predicted the secondary structure with the lowest free energy of the NTO3 ssRNA (**Supplementary Figure 1 a and b**) and of the NTO3-ORF62-5’ dsRNA hybrid, using both the unedited and edited forms of the asRNA (**Supplementary Figure 1 c and d**). When forming an intramolecular secondary structure, the unedited form has a higher free energy state than the edited form (−143.2 kcal/mol compared to −169.4 kcal/mol), which suggests that hyper-editing confers thermodynamic stability to the secondary structure of the RNA. Twelve of the 17 Is may aid in the formation of stem structures, while in the unedited RNA only four of the As in the same position have a complementing nucleotide. Contrarily, the secondary structure of the dsRNA formed by the edited NTO3 and the ORF62 is slightly less stable than that formed by the unedited form (−818.7 kcal/mol compared to −822.9 kcal/mol), but allows the formation of identical secondary structures.

**Figure 9.**
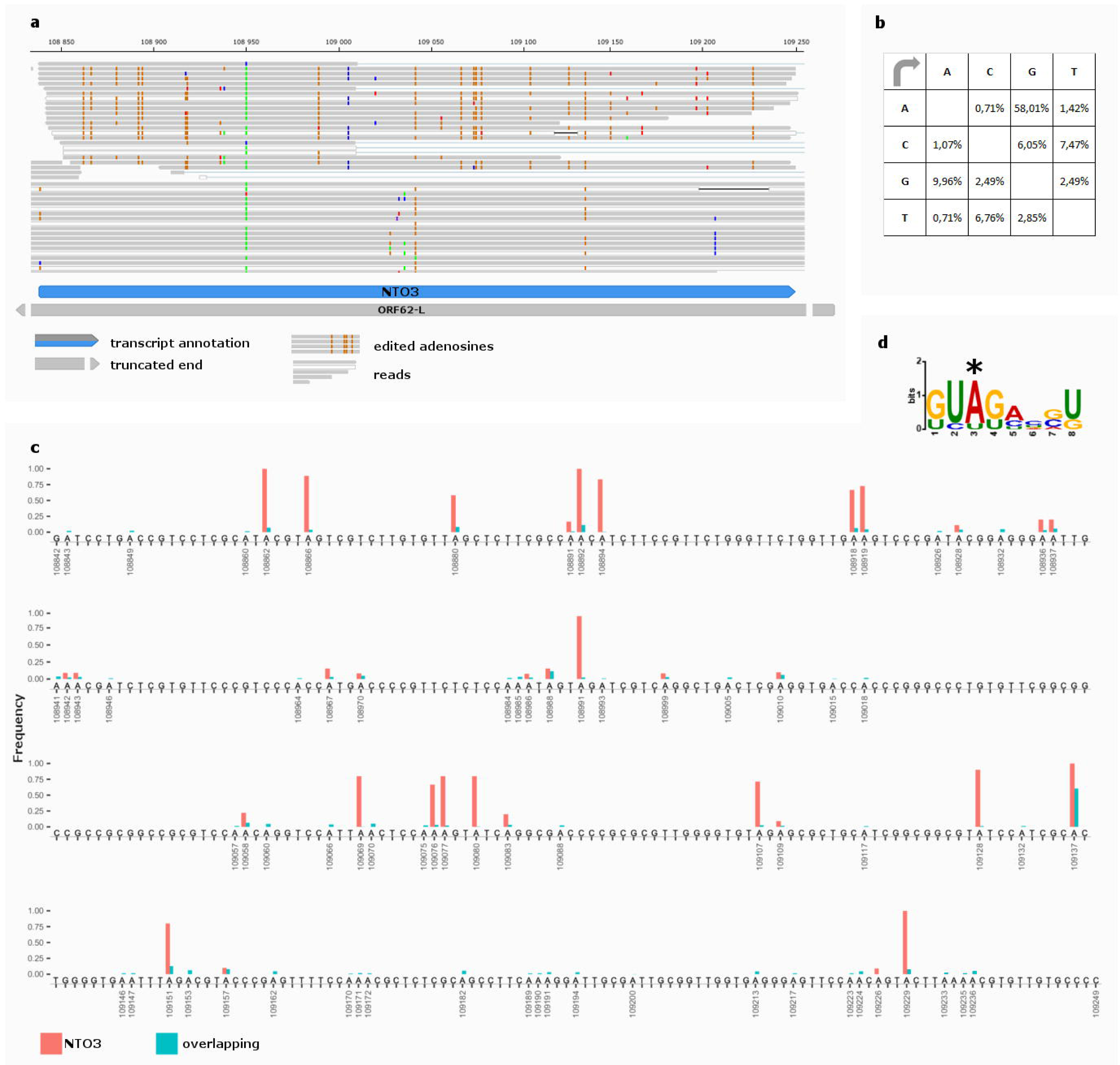
A to I hyper-editing of NTO3. a. Reads of NTO2 and the overlapping transcripts mapping to the VZV genome visualized with IGV. The orange dots represent G mismatches, indicating editing events. b. Substitution matrix of the NTO2 reads (n=703 substitutions / 7749 nt, p<0.0001, Fisher’s exact test). c. The position and frequency of A->G substitutions on the genomic sequence corresponding to the NTO2 transcript, showing both NTO2 and the overlapping transcripts. Substitutions with high frequencies indicate A to I editing events, while those with low frequency are sequencing errors. d. The motif surrounding the editing sites. The motif was found using the MEME software suit with an E-value of 0.58 and a log likelihood ratio of 46. The edited adenosine is marked with a *.

## Discussion

Until now, the VZV transcriptome have been analyzed by Northern blot, primer extension, microarray and Illumina sequencing [17, 68–75]. These techniques have generated useful data, but they have limitations to provide a comprehensive list of the VZV transcripts. In this work, we used the ONT MinION LRS technique for the investigation of the poly(A)+ fraction of the VZV transcriptome. Our results identified altogether 151 novel transcripts, including novel mRNA molecules, monocistronic transcripts, transcript isoforms, as well as multigenic transcripts. Novel splice sites and splice variants were also detected. In this work, we discovered 5 sncRNAs and 23 lncRNAs and annotated their TSSs and TESs with base pair precision.

The enrichment of As and Us upstream of PASs in mammalian systems is well-established [76–78]. These homopolymer A stretches may also cause false priming and template switching events during reverse transcription, which results in false TESs. In our work, we excluded these artefacts by the use of our analysis pipeline, and demonstrated the verity of the annotated TESs by the presence of the canonical GU-rich region in the +10 interval downstream of TESs [79].

Similar to other herpesviruses [21, 50], this study also revealed a complex meshwork of transcriptional read-throughs and overlaps. It has been earlier known that the polycistronic transcription units include co-terminal multigenic RNA molecules, which represent a large extent of parallel overlaps along the entire viral genome. According to our current knowledge, the downstream genes on these long transcripts are untranslated. We can raise the question as to whether the untranslated downstream sequences have any function or if they are mere random read-through products representing transcriptional noise. We can address the same question for the convergently and divergently overlapping RNA segments, and also for the alternative transcriptional overlaps. We have put forward a hypothesis that explains the potential role of this phenomenon which is based on transcriptional interaction between the RNA polymerase molecules at the overlapping regions. Since essentially every gene produces overlapping transcripts and therefore the pairwise interactions can spread along the genome thereby forming a transcriptional interference network (TIN), which alongside the promoter-transcription factor system determines the global gene expression pattern of the viral DNA [51]. It has been previously shown that orf63 has no trans-regulatory effect on the expression of orf62 [80], however deletion of orf63 increases the expression of orf62 [81], suggesting that there is a link between the regulation of the two genes. A possible explanation could be the transcriptional interference caused by the head-to-head overlaps between ORF62 and NTO1v1 and NTO1v2.

Polycistronic (especially bicistronic) transcripts are also common in eukaryotic organisms, but their generation is explained by trans-splicing and not by transcriptional read-through [82]. However, unprocessed transcripts are difficult to detect because of their short existence, therefore, it cannot be excluded that these transcripts are the result of a transcription read-through mechanism. The predominant occurrence of adjacent genes in the chimeric transcripts suggests that transcriptional readthrough followed by cis-splicing may be the case, that is, the existence of a large extent of transcriptional read-throughs may be not restricted to the herpesviruses but they may represent a general phenomenon. Furthermore, the antisense RNAs produced from their own promoters or by transcriptional read-throughs may hybridize with their sense counterparts thereby initiating RNA interference [83]. Very long complex transcripts form a distinct category among multigenic transcripts because of their oppositely oriented ORFs and increased size. Similarly to other herpesviruses [21, 50] they are present in very low abundance in VZV, however their existence is ambiguous, as they can be formed during reverse transcription by template switching [84]. It has been previously thought that the OriS of VZV lacks any overlapping transcripts [6]. Additionally, Davison and colleagues (1986) hypothesis the existence of non-coding RNA in the intergenic region between orf62 and orf63. We detected several transcripts overlapping OriS, and a small non-coding RNA (the NCO3) between the before-mentioned two genes. This region of the viral transcriptome is structurally similar to the PRV transcriptome. In the vicinity of OriS, we identified two additional novel non-coding transcripts (NTO2 and NTO4) and two 5’ elongated and spliced version of ORF63. These nroRNAs are supposed to play a role in the regulation of the viral replication [85] through the interplay between the transcriptional and replication apparatuses [51]. Specifically, the role of the transcription of these RNA molecules might be to interfere with the replication machinery, in order to force the replication fork to an unidirectional progression [63].

Additionally, we detected a hyper-editing process in the novel NTO3 transcript. This phenomenon was previously observed in other members of the herpesvirus family [86–88], but to our knowledge, this is the first observation of A to I hyper-editing in alphaherpesviruses. This process plays a crucial role in the cell’s innate immunity [89], while it can be hijacked by some viruses to evade inactivation [29]. Additionally, in hepatitis delta virus hyper-editing is indispensable for replication [30]. A to I editing decreases the affinity of the antisense transcript to the sense RNA, destabilizing their interaction, which may affect the binding of dsRNA enzymes like RNase III homologues [90]. Our *in-silico* analysis shows that despite elevating the level of free energy of the sense-antisense RNA hybrid, hyper-editing does not result in a change of the secondary structure, but in the formation of I·U wobble pairs which are significantly more resistant to possible Dicer cleavage [91], inhibiting RNA interference. It is also possible that I·U pairs play a role in the cleavage and degradation of ORF62 by a process mediated by the Tudor staphylococcal nuclease (Tudor-SN) [92, 93]. Further investigation is needed to elucidate the significance of hyper-editing on the NTO3. An evaluation of gene expression at different time points could shed light on the presence of the asRNA and on the amount of editing in different stages of the viral life cycle. Additionally, miRNA assays of the viral transcriptome in different time points of the infection could prove the effect of the RNAi on the dsRNA formed by the sense-antisense pair.

Splice events are thought to be rare in alphaherpesviruses, however, they appear to be underestimated in the light of LRS techniques. In this work, we enriched the list of spliced transcripts of the lytic phase of the viral infection, and we identified the novel combinations of splice sites in ORF42/45.

At least six VZV-transcripts have been shown to be expressed in latency [94, 95], however recent target enrichment SRS data suggests that VLT and ORF63 are the only two expressed transcripts, maintaining the dormant phase of VZV. In this work Depledge and coworkers showed that the VLT is spliced differently in the latent and lytic phase of the viral lifecycle, and identified several length and splice isoforms during productive infection [9]. We confirmed the existence of four introns of this transcript in the longer, lytic forms of VLT. Additionally, we showed a splice isoform of ORF63 and ORF64 possessing the same introns as the VLT. This suggests that occasionally VLT, ORF63 and ORF64 can form a single transcriptional unit. We found that a novel splice isoform of ORF63 produces a long transcript, which has non-canonical splice donor site (GC, instead of GU). Another rare but clearly visible splice variant of ORF63-L has the same, non-canonical splice donor site. The reason for this could be simply the higher GC content of these regions. Another explanation may be that the splice isoforms of ORF63 are recognized by host cell spliceosomes differentially, in order to perform varying expression pattern during the life cycle of virus. Similar alternative splicing has been described in VLT, assuming distinct functions for it in lytic and latent phase.

The uORFs are supposed to play a regulatory role in the translation of eukaryotic mRNAs. Our results suggests, that at least some of the uORFs could play a role in the production of N-terminally truncated proteins, while most of them have a large-enough space between their stop codons and the protein coding ORF’s AUG for a reinitiation event, thus resulting in unaltered protein translation [96]. Further studies implying ribosome profiling and mRNA and protein expression studies are needed to determine the precise function of these uORFs. Nevertheless, the use of alternative promoters for producing TSS variants with or without uORFs may have a role in providing a differential control of translation at distinct stage of the viral lifecycle [21].

## Conclusions

This study substantially redefines the VZV transcriptome by identifying a large number of novel RNA molecules and transcript isoforms, as well as revealing a complex pattern of transcriptional overlaps. The extensive transcriptional overlaps may indicate an interaction between the transcription machineries. Additionally, the discovery of nroRNAs suggests an interference between the replication and transcription apparatuses. Besides the significant advance in transcriptome annotation, these data may also help in controlling this virus.

## List of abbreviations

ADAR1: adenosine deaminase acting on RNA type 1
AS: antisense
AT: alternative termination
HSV-1: herpes simplex virus type 1
HCMV: human cytomegalovirus
lncRNA: long non-coding RNA
LRS: long-read sequencing
ncRNA: non-coding RNA
nroRNA: near-replication-origin RNA
NTO: near to (replication) origin
ONT: Oxford Nanopore Technologies
PAS: Poly(A) signal
PRV: pseudorabies virus
RT: reverse transcriptase
sncRNA: short non-coding RNA
SRS: short-read sequencing
TES: transcription end site
TR: truncated
TSS: transcription start site
uORF: upstream open reading frame
UTR: untranslated region
VLT: VZV latency transcript
VLT_ly_: lytic form of VLT
VZV: Varicella Zoster Virus

## Ethics approval and consent to participate

Not applicable.

## Consent for publication

Not applicable.

## Availability of data and material

The sequencing data and the transcriptome assembly have been uploaded to the European Nucleotide Archive under the project accession number PRJEB25401.

## Competing interests

The authors declare that they have no competing interests

## Funding

This work was supported by the Swiss-Hungarian Cooperation Programme [SH/7/2/8] to ZBo. The work was also supported by the Bolyai János Scholarship of the Hungarian Academy of Sciences to DT.

## Author Contributions

IP analyzed the data, participated in RNA-isolation and sequence alignment and drafted and wrote the manuscript. NM participated in data analysis, carried out the statistical analysis, prepared the figures and revised the manuscript. DT carried out the ONT MinION cDNA sequencing, participated in the design of the study, and took part in drafting the manuscript. AS participated in the sequence alignment and carried out the *in silico* analysis. ZC prepared the RNA, DNA and cDNA samples, carried out the PCR analysis and participated in the MinION sequencing. KM carried out the virus infection and propagated the cells. ZBo conceived, designed and coordinated the study and wrote the manuscript. All authors have read and approved the final version of the manuscript.

---

**Supplementary Figure 1 The secondary structure of NTO3 (a. and b.) as well as the hybrid formed by the ORF62-5’ fragment and the NTO3 (c and d).**

a. The secondary structure of the unedited ORF62-AS2 with a free energy of −143.2 kcal/mol. The adenines in the editing sites are colored in green.

b. The secondary structure of the edited NTO3 with a free energy of −169.4 kcal/mol. The inosines in the editing sites are colored in orange.

c. The secondary structure of the sense-antisense hybrid composed of the first 467 bases of ORF62 labeled ORF62-5’ (gray) and the full sequence of NTO3 (blue), the latter is its unedited form. The free energy of the structure formed by the two molecules is −822.9 kcal/mol.

d. The secondary structure of the sense-antisense hybrid composed of the first 467 bases of ORF62 labeled ORF62-5’ (gray) and the full sequence of NTO3 (blue), the later in its edited form. The free energy of the structure formed by the two molecules is −818.7 kcal/mol. The position of I·U base pairs is marked with black arrows.

**Additional File Legend**

The additional file is in .xlsx format, containing eight Supplementary Tables on separate pages and a References page.

Supplementary Table 1. The list of transcripts with TSS and TES that passed our criteria for validation.

Supplementary Table 2. The list of transcripts with 5’ and 3’ ends that did not pass our criteria for validation.

Supplementary Table 3. Promoter motifs and poly(A) signals.

Supplementary Table 4. Overlaps between the transcripts listed in Supplementary Table 1.

Supplementary Table 5. The full list of 5’ ends in the non-cap-selected datasets generated by our pipeline. The p-values were calculated using the Poisson distribution. Positions excluded because of missing TESs are marked by *, while those excluded because of false priming or strand switching are marked with ** in the Notes column. The number of 5’ ends in the ±50 vicinity of a given position are shown in the Vicinity column.

Supplementary Table 6. The full list of 5’ ends in the cap-selected datasets generated by our pipeline. The p-values were calculated using the Poisson distribution. Positions excluded because of missing TESs are marked by *, while those excluded because of false priming or strand switching are marked with ** in the Notes column. The number of 5’ ends in the ±50 vicinity of a given position are shown in the Vicinity column.

Supplementary Table 7. The full list of 3’ ends in the non-cap-selected datasets generated by our pipeline. The p-values were calculated using the Poisson distribution. Positions excluded because of missing TSSs are marked by *, positions excluded because of false priming or strand switching are marked with **. TES with more than 3 but less than 5 adenines in the vicinity of the TES with *y and they were accepted. The number of 3’ ends in the ±50 vicinity of a given position are shown in the Vicinity column.

Supplementary Table 8. The full list of 3’ ends in the cap-selected datasets generated by our pipeline. The p-values were calculated using the Poisson distribution. Positions excluded because of missing TSSs are marked by *, positions excluded because of false priming or strand switching are marked with **. TES with more than 3 but less than 5 adenines in the vicinity of the TES with *y and they were accepted. The number of 3’ ends in the ±50 vicinity of a given position are shown in the Vicinity column.

